# The CXCL12/CXCR4 signalling axis retains neutrophils at inflammatory sites in zebrafish

**DOI:** 10.1101/626978

**Authors:** Hannah M. Isles, Kimberly Herman, Anne L. Robertson, Catherine A. Loynes, Lynne R. Prince, Philip M. Elks, Stephen A. Renshaw

**Author notes:** **Corresponding Authors:** Stephen A. Renshaw, Philip M. Elks, The Bateson Centre, University of Sheffield, Firth Court, Western Bank, Sheffield S10 2TN, UK.

## Abstract

The inappropriate retention of neutrophils in the lung is a major driver of the excessive tissue damage characteristic of respiratory inflammatory diseases including COPD, ARDS and cystic fibrosis. The molecular programmes which orchestrate neutrophil recruitment to inflammatory sites through chemotactic guidance have been well studied. However, how neutrophil sensitivity to these cues is modulated during inflammation resolution is not understood. The identification of neutrophil reverse migration as a mechanism of inflammation resolution and the ability to modulate this therapeutically has identified a new target to treat inflammatory disease. Here we investigate the role of the CXCL12/CXCR4 signalling axis in modulating neutrophil retention at inflammatory sites. We used an *in vivo* tissue injury model to study inflammation using transgenic zebrafish larvae. Expression of *cxcl12a* and *cxcr4b* during the tissue damage response was assessed using *in situ* hybridisation and analysis of RNA sequencing data. CRISPR/Cas9 was used to knockdown *cxcl12a* and *cxcr4b* in zebrafish larvae. The CXCR4 antagonist AMD3100 was used to block the Cxcl12/Cxcr4 signalling axis pharmacologically. We identified that *cxcr4b* and *cxcl12a* are expressed at the wound site in zebrafish larvae during the inflammatory response. Following tail-fin transection, removal of neutrophils from inflammatory sites is significantly increased in *cxcr4b* and *cxcl12a* CRISPR knockdown larvae. Pharmacological inhibition of the Cxcl12/Cxcr4 signalling axis accelerates inflammation resolution, an effect caused by an increase in neutrophil reverse migration. The findings of this study suggest that CXCR4/CXCL12 signalling may play an important role in neutrophil retention at inflammatory sites, identifying a potential new target for the therapeutic removal of neutrophils from the lung in chronic inflammatory disease.

## Introduction

The inappropriate retention of activated innate inflammatory cells at inflammatory sites is major driver of chronic inflammatory diseases including asthma, COPD and rheumatoid arthritis [1]. Neutrophils are the first cells recruited to the site of an inflammatory stimulus, where they are potent anti-microbial effectors through the phagocytosis of foreign material, generation of reactive oxygen species and the production of extracellular traps [2]–[4]. These non-specific anti-microbial mechanisms promote a tissue microenvironment which is unfavourable to pathogens, but at the expense of host tissue integrity [5]. Neutrophil removal from inflammatory sites is therefore tightly regulated to minimise collateral tissue damage, thereby preventing chronic inflammatory disease [6]. Despite the global burden of chronic inflammatory diseases, there are currently no effective therapies to treat the neutrophilic component of these conditions, highlighting a need to identify novel drug targets to promote the successful resolution of inflammation.

It has been known for thirty years that neutrophils undergo apoptosis followed by efferocytosis by macrophages, and this is the best characterised mechanism by which neutrophils are removed from inflammatory sites [7], [8]. Although methods to both accelerate and delay apoptosis have been identified [9]–[13], none of these are yet in clinical use for inflammatory disease. More recently, reverse migration has been identified as a mechanism by which neutrophils redistribute into the tissue or vasculature surrounding the inflammatory site, an anti-inflammatory mechanism which is thought to disperse the inflammatory burden [12]–[15]. The mechanisms governing this newer phenomenon are not fully understood, though it is clear that the capacity of neutrophils to cause host tissue damage is increased when either apoptosis or reverse migration are impaired, resulting in the inappropriate retention of neutrophils at the inflammatory site [16]. Understanding neutrophil reverse migration represents novel therapeutic avenues to treat neutrophil mediated chronic inflammation.

During inflammation, neutrophils respond to complex guidance cues provided in part by chemokine gradients which promote the directed migration of neutrophils from the circulation and into inflamed tissues [17]. More recently, a role for chemokine signalling in modulating neutrophil reverse migration has been identified [15], [18], making chemokine receptors an attractive target for investigation. Computational modelling and *in vivo* studies of reverse migration have shown that this process likely occurs as a result of the stochastic redistribution of neutrophils following their desensitisation to local chemotactic gradients over time [12], [18], [19]. In zebrafish, neutrophil reverse migration can be delayed by stabilising HIF1α which promotes neutrophil retention at inflammatory sites [13], suggesting that downstream HIF signalling targets retain neutrophils at inflammatory sites. Work by our group and others has shown that this retention of neutrophils at inflammatory sites is both mechanistically important [13], [16], and can be manipulated therapeutically [10], [12], [18], yet the molecular mechanisms remain to be elucidated.

CXCR4 is a G protein coupled receptor expressed by many leukocytes, which exerts its biological functions by signalling through its major ligand CXCL12 (formerly known as stromal derived factor 1). CXCL12/CXCR4 signalling is a key retention signal for neutrophil release into the blood circulation from hematopoietic tissues, the crucial role of which is highlighted in patients with warts, hypogammaglobulinaemia, infection and myelokathesis (WHIM) syndrome. Gain of function WHIM mutations result in increased CXCR4 signalling, the consequence of which is severe neutropenia with increased neutrophil retention in the bone marrow [20].

There is growing evidence to support a role for CXCL12/CXCR4 in neutrophil retention in the context of inflammatory disease. Tissue infiltrated neutrophils from patients with chronic inflammatory lung diseases and rheumatoid arthritis have increased CXCR4 surface expression [21]. Neutrophil surface expression of CXCR4 is increased after extravasation into injured lungs in mice [22] and in human tissue samples, where pulmonary CXCL12 expression increases during acute lung injury [23]. Additionally, the inhibition of CXCL12 using blocking antibodies prevented the accumulation of neutrophils in the lung during the late stages of LPS induced lung injury [22]. Based on this evidence we hypothesised that CXCL12/CXCR4 functions as a retention signal in the context of tissue damage, functioning to maintain active neutrophils at the inflammatory site.

Here we present a new role for the CXCL12/CXCR4 signalling axis in the retention of neutrophils at inflammatory sites and demonstrate a role for neutrophil retention signalling in modulating inflammation resolution in zebrafish larvae. Using both pharmacological and genetic approaches to manipulate the CXCL12/CXCR4 signalling axis, we demonstrate that interruption of CXCR4 signalling accelerates inflammation resolution by increasing neutrophil reverse migration. We have identified a druggable target which could be a therapeutic target to remove inappropriately retained neutrophils from inflammatory sites during disease.

## Results

### *cxcr4b* and *cxcl12a* are expressed following tissue damage in zebrafish

Zebrafish have two paralogues for CXCR4 and CXCL12, following a genome duplication event in teleost evolution. The expression of *cxcr4a* and *cxcr4b* is mutually exclusive in most cell lineages, indicating roles in different tissues. [24]. To determine the gene expression of Cxcr4 and Cxcl12 during the cellular response to tissue damage in zebrafish larvae, we first investigated neutrophil expression of *cxcr4 and cxcl12*. We studied published datasets combining RNA sequencing of zebrafish larval neutrophils and single-cell RNA sequencing data from adult zebrafish blood lineages [25], [26]. In adult zebrafish neutrophils, *cxcr4b* is highly expressed by the neutrophil lineage whilst *cxcr4a* is undetectable (Figure 1A-B). *Cxcl12a* is expressed by a small population of adult zebrafish neutrophils, albeit far fewer than *cxcr4b*, whilst *cxcl12b* is expressed by very few cells (Figure 1C-D). We analysed larval stage neutrophil RNA sequencing data [25], and found that fragments per kilobase million (fpkm) values for *cxcr4b* were over 100-fold higher than the fpkm values for *cxcr4a* (Figure 1E), confirming that *cxcr4b* is the predominantly expressed isoform in larval zebrafish neutrophils. Furthermore, we confirmed that expression of *cxcl12a* and *cxcl12b* was low in larval neutrophils (Figure 1F).

**Figure 1.**
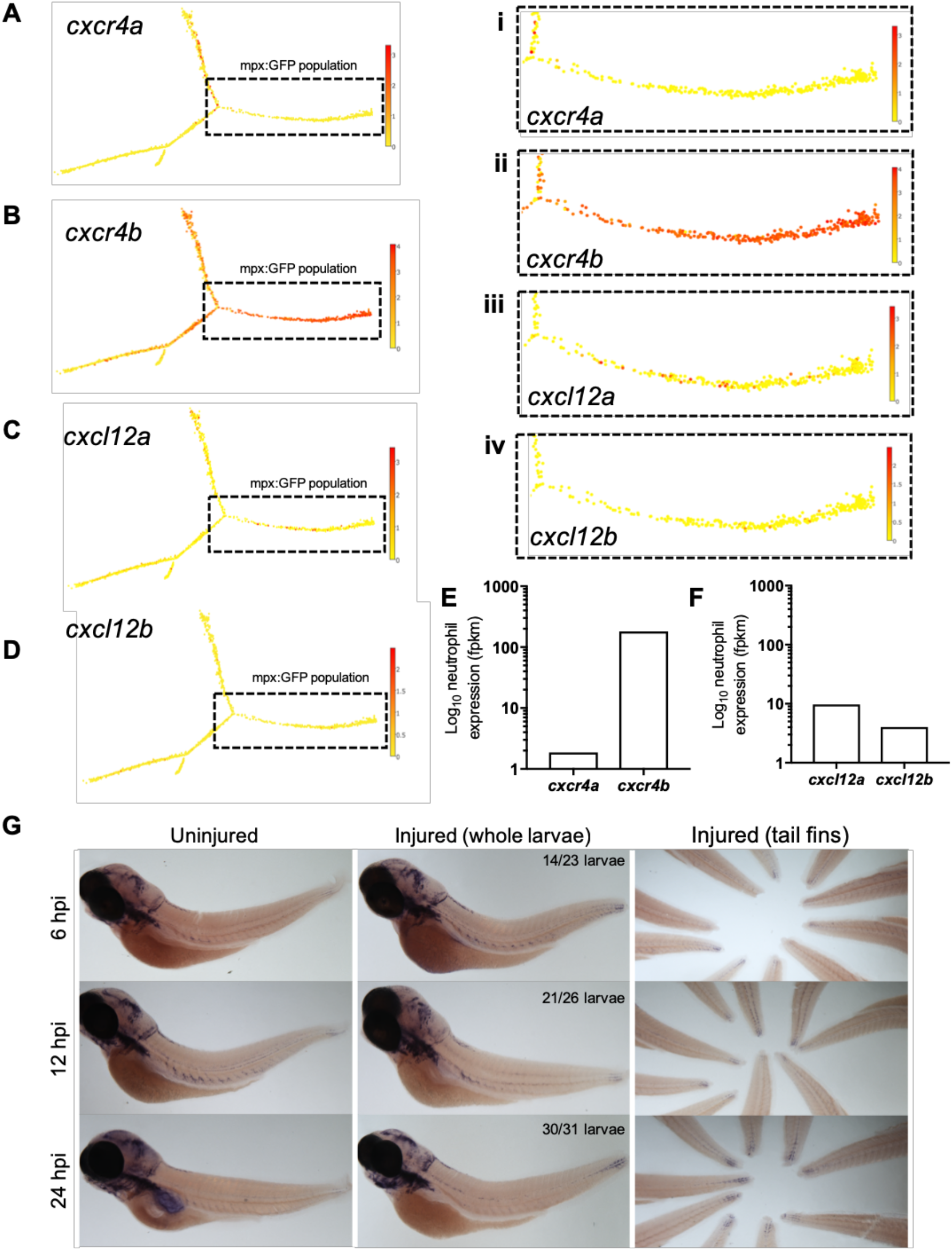
*cxcr4b* and *cxcl12a* are expressed following tissue damage in zebrafish. **A-D** Single-cell gene expression profiles for *cxcr4* and *cxcl12* in the zebrafish blood lineage. Single cell gene expression values extracted from the Sanger BASiCz zebrafish blood atlas. Circles represent individual cells colour coded where red is high expression and yellow is no expression. Neutrophil lineage (mpx:GFP positive) is highlighted by black dashed box and expanded in (i-iv). **E-F** RNA sequencing of FACS sorted GFP positive cells from *TgBAC(mpx:GFP)i114* zebrafish larvae at 5 days post fertilization. FPKM values illustrate neutrophil expression of (E) *cxcr4a* and *cxcr4b* and (F) *cxcl12a* and *cxcl12b.* **G** Whole mount *in situ* hybridization using an antisense DIG labelled RNA probe for *cxcl12a* mRNA. Wildtype *nacre* zebrafish larvae were injured and fixed in PFA at 6, 12 and 24 hours post injury, along with uninjured age-matched control larvae. Left and middle panels show whole zebrafish larvae at timepoints indicated, right panel shows tailfins of a representative experiment. Quantification shows number of larvae which look like representative image from 2 independent experiments.

Zebrafish Cxcr4b is activated by the chemokine Cxcl12a [27], hence we investigated the expression of *cxcl12a* during the inflammatory response. To induce an inflammatory response we used our well characterised tail-fin injury model of spontaneously-resolving inflammation[28], where neutrophil recruitment is observed between 0-6 hours post injury (hpi) and inflammation resolution occurs between 6-24hpi. Whole mount *in situ* hybridisation was used to detect *cxcl12a* mRNA at the wound site in 3dpf larvae following tail fin transection. *Cxcl12a* mRNA expression was detected in injured larvae as early as 6hpi during the recruitment phase (Figure 1G). Interestingly, *cxcl12a* mRNA expression continued to increase throughout the resolution phase up to 24hpi (Figure 1G) in keeping with other reports of *cxcl12* expression following fin injury. These findings show the expression of *cxcr4b* by neutrophils and *cxcl12a* at the tissue injury site during the inflammatory response in zebrafish.

### Genetic manipulation of the CXCL12/CXCR4 signalling axis accelerates inflammation resolution

After determining that *cxcl12a* was expressed at the wound site in injured larvae, we next investigated neutrophil responses to tissue injury in the absence of the CXCL12/CXCR4 signalling axis. We hypothesised that if CXCL12/CXCR4 signalling was a neutrophil retention signal, inhibition of this pathway would accelerate inflammation resolution. We used CRISPR/Cas9 to study the role of Cxcl12a and Cxcr4b in neutrophilic inflammation resolution using the *TgBAC(mpx:GFP)i114* transgenic reporter line [28]. A crRNA targeting the pigment gene tyrosinase (*tyr*) [29] was used for control injections and to allow for visual identification of successful knockdown. Knockdown of *tyr* produces an albino phenotype in zebrafish larvae (Supplemental Figure 1A-B) without affecting neutrophil development or the neutrophilic inflammatory response (Supplemental Figure 1C-D). We generated *cxcr4b* or *cxcl12a* ‘crispants’ (newly generated “F0” CRISPR/Cas9-mediated mutants) and transected tail-fins at 2 dpf, counting neutrophils at the wound site at 4, 8 and 24 hpi (Figure 2A). Neutrophil counts in *cxcr4b* crispants were significantly increased at the wound site during the neutrophil recruitment phase (4hpi), consistent with enhanced release of *cxcr4b* mutant neutrophils from their site of production [30] (Figure 2B). *Cxcl12a* crispants showed no difference in neutrophil recruitment (Figure 2B). No significant difference in neutrophil numbers at the wound site was detected between groups at 8 and 24hpi (Figure 2B). To control for the increase in early neutrophil recruitment measured in Cxcr4b crispants, we calculated percentage inflammation resolution scores in individual larvae between 4 and 8 hpi. Both Cxcr4b and Cxcl12a crispants had significantly higher percentage inflammation resolution compared to control larvae (Figure 2C). Whole body neutrophil numbers were not affected in *cxcr4b* crispants, but were significantly reduced in *cxcl12a* crispants (Figure 2D). These data demonstrate that loss of Cxcl12/Cxcr4 signalling accelerates inflammation resolution in zebrafish larvae, suggesting that the CXCL12/CXCR4 signalling axis is required for neutrophil retention at inflammatory sites.

**Figure 2.**
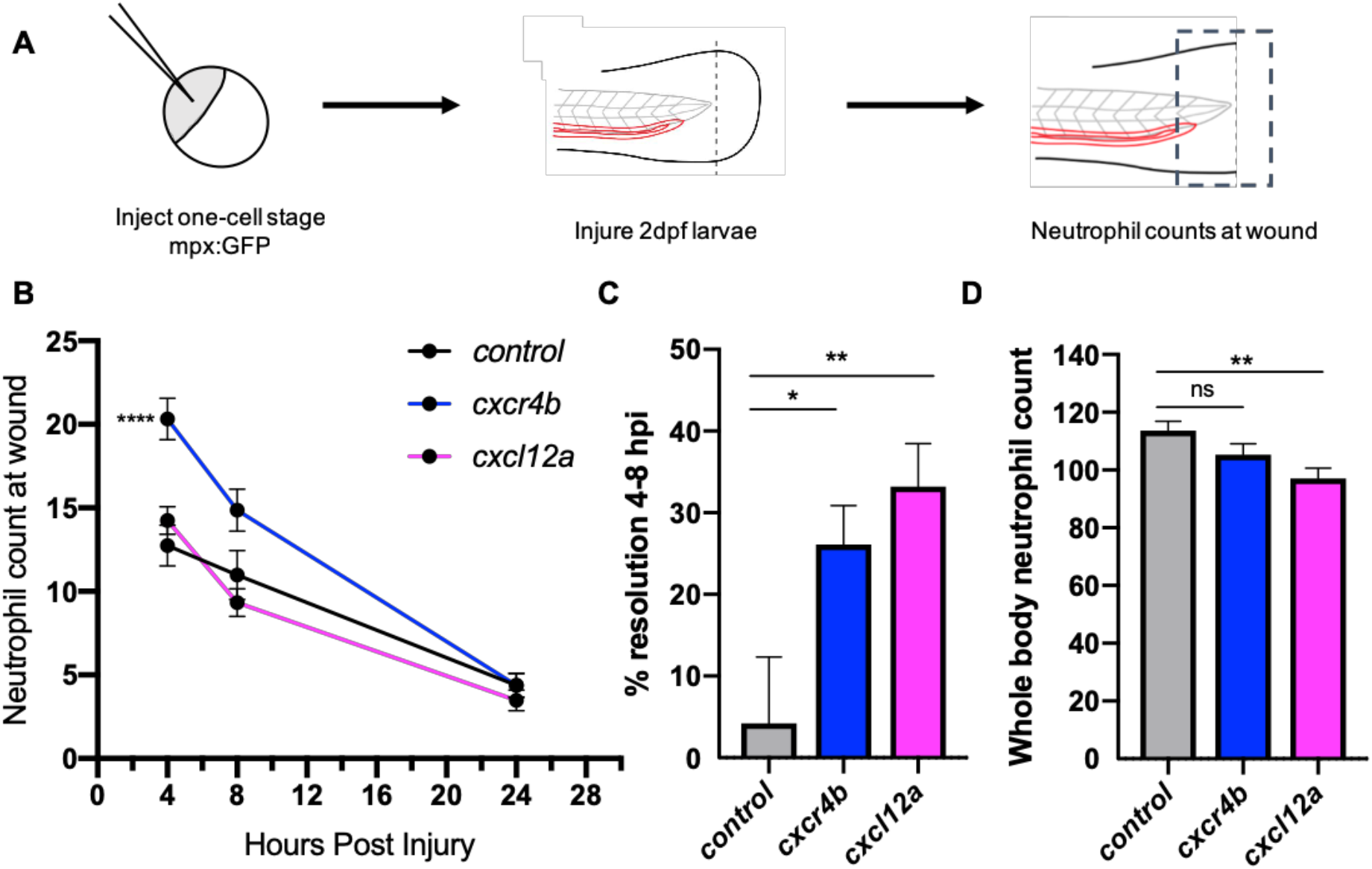
Knockdown of *cxcr4b* using CRISPR/Cas9 accelerates inflammation resolution. **A** Experimental schematic of CRISPR/Cas9 experiments in 2dpf mpx:GFP larvae. **B** CRISPR/Cas9-mediated knockdown of *cxcr4b* and *cxcl12a* accelerates inflammation resolution. Neutrophil counts at the wound site in control *tyr* crRNA injected larvae (black line), *cxcr4b* crRNA injected larvae (blue line), and *cxcl12a* crRNA injected larvae (pink line) at 4, 8 and 24hpi. Error bars shown are mean ± SEM. Groups were analysed using an ordinary one-way ANOVA and adjusted using Tukey’s multi comparison test. ****p<0.001 n=36 from 3 independent experiments. **C** % inflammation resolution was calculated between 4-8hpi. Groups were analysed using an ordinary one-way ANOVA and adjusted using Tukey’s multi comparison test. *p<0.04, **p<0.004. **D** Whole body neutrophil numbers were measured in mpx:GFP crispant larvae at 2dpf. n=30-35 per group from 3 independent experiments. Error bars shown are mean ± SEM. Groups were analysed using an ordinary one-way ANOVA and adjusted using Tukey’s multi comparison test, **p<0.005.

### Pharmacological inhibition of CXCR4 accelerates inflammation resolution

Genetic knockdown of CXCR4 signalling causes neutrophil release from the caudal haematopoietic tissue (CHT), enhancing neutrophil recruitment, confounding assessment of inflammation resolution. To circumvent this, we used the CXCR4 antagonist AMD3100 to block CXCR4 signalling in a time-sensitive fashion (Figure 3A). At 8hpi a significant decrease in neutrophil counts at the wound site was detected in AMD3100 treated larvae (Figure 3B). Percentage inflammation resolution was significantly higher in AMD3100 treated larvae (Figure 3C), whilst whole body neutrophil counts were not affected by AMD3100 at 24 hours post administration (Figure 3D). Together these data demonstrate that pharmacological inhibition of CXCR4 in larvae which have mounted a normal response accelerates inflammation resolution, further supporting a role for CXCL12/CXCR4 signalling in neutrophil retention signalling at sites of tissue damage.

**Figure 3.**
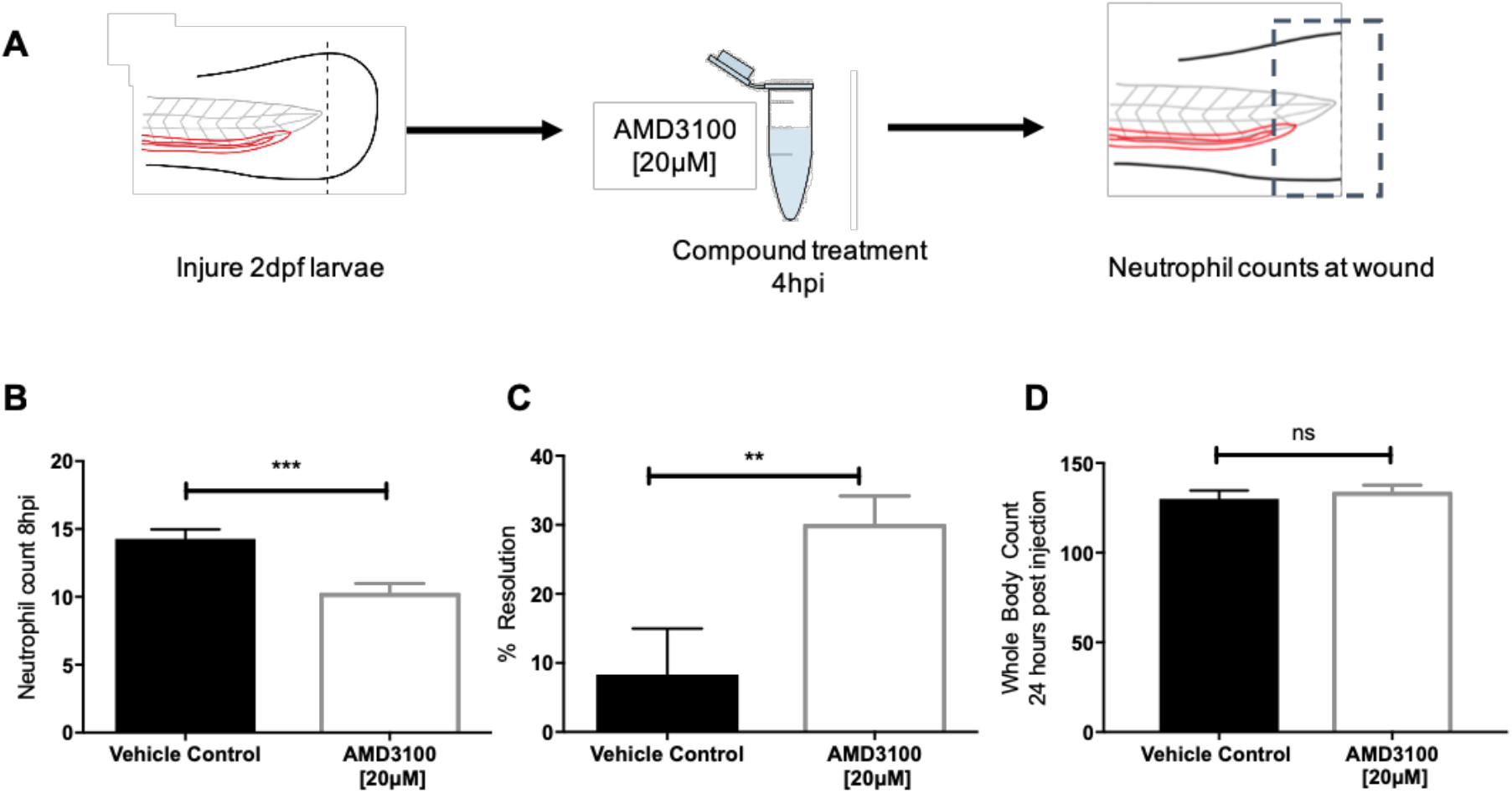
Inhibition of CXCR4 using AMD3100 accelerates inflammation resolution. **A** Experimental schematic of inflammation resolution experiments with AMD3100 compound treatment. **B** Number of neutrophils at the wound site in injured 2dpf mpx:GFP larvae treated with AMD3100 or vehicle control at 8hpi. Groups were analysed using an unpaired t-test, ***p<0.0002, n=55 larvae from 5 independent experiments. **C** % inflammation resolution for larvae treated with vehicle control or AMD3100. Groups were analysed using an unpaired t-test, **p<0.008 n=32 larvae from 3 independent experiments. **D** Whole body neutrophil counts in 3dpf mpx:GFP larvae 24 hours post administration of AMD3100 or vehicle control. Groups were analysed using an unpaired t-test, n=26 larvae from 3 independent experiments.

### Inhibition of CXCL12/CXCR4 signalling increases neutrophil reverse migration

Two principal mechanisms of inflammation resolution have been described: neutrophil apoptosis followed by efferocytosis by macrophages and reverse migration of neutrophils away from inflammatory sites. We have previously proposed that neutrophil release from inflammatory sites is best explained by the desensitisation of neutrophils to local chemokine gradients [19]. This led us to the specific hypothesis that inhibition of CXCL12/CXCR4 signalling would accelerate reverse migration by accelerating neutrophil desensitisation to CXCL12 gradients. To study neutrophil reverse migration, we used a well described photoconversion approach to study the reverse migration of neutrophils from a wound site [10], [12], [13], [31]. AMD3100 was administered to *TgBAC(mpx:GAL4-VP16); Tg(UAS:Kaede)i222* (referred to as mpx:kaede) larvae at 5hpi and neutrophils at the wound site were photoconverted and tracked during the resolution phase (Figure 4A). Neutrophil migration away from the wound site was significantly higher in larvae treated with AMD3100 (Figure 4B), an effect which was not due to a difference in the number of photoconverted neutrophils (Figure 4C). Together these data demonstrate that inhibition of CXCL12/CXCR4 signalling can increase inflammation resolution by accelerating neutrophil reverse migration, identifying this signalling axis as a potential therapeutic target to specifically remove inflammatory neutrophils without affecting the normal recruitment of neutrophils to new inflammatory or infectious lesions.

**Figure 4.**
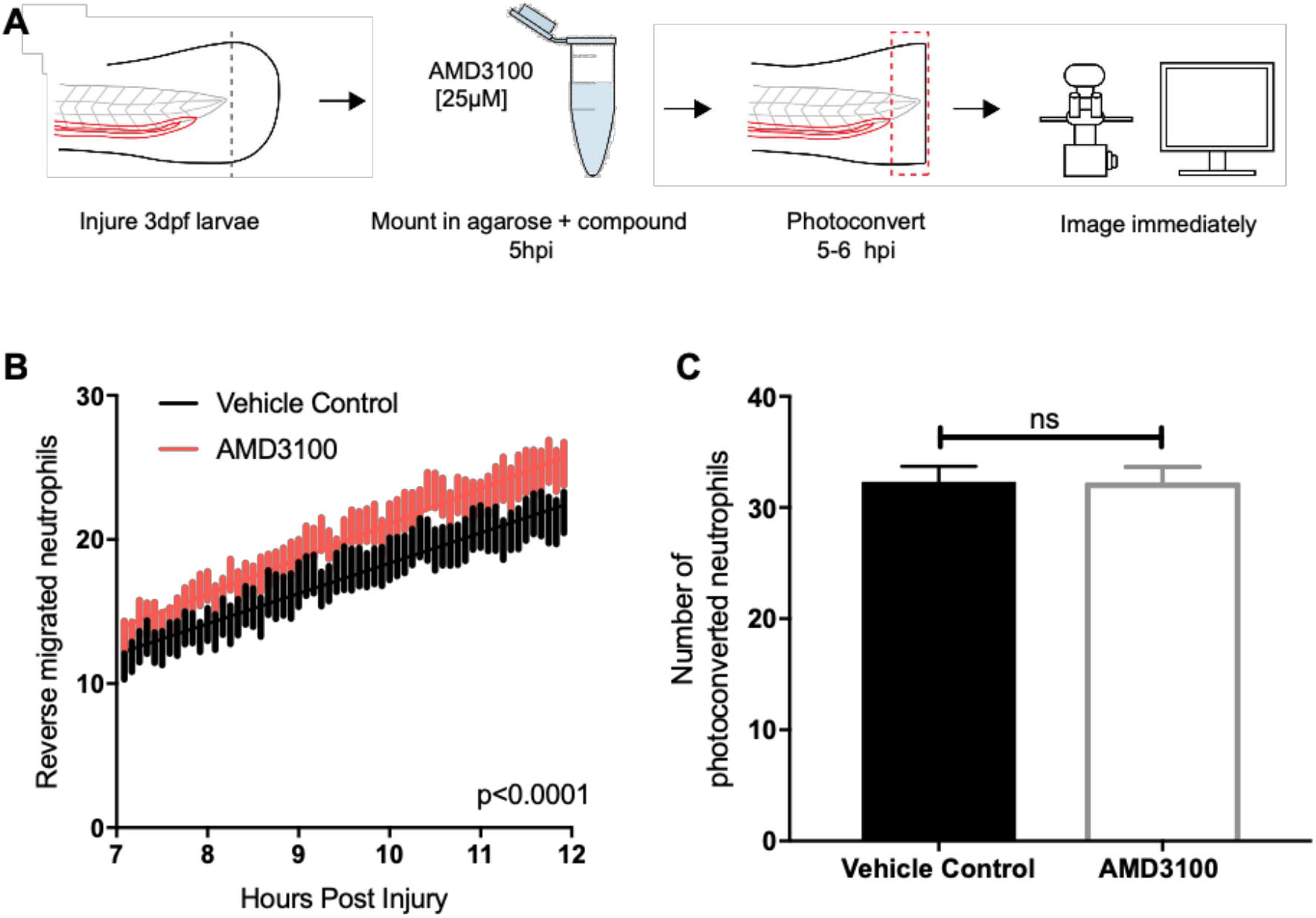
Inhibition of CXCR4 using AMD3100 accelerates neutrophil reverse migration. **A** Experimental schematic of neutrophil reverse migration assay. Tail fin transection was performed on 3dpf mpx:kaede larvae. Larvae were mounted in a 1% agarose solution containing AMD3100 or vehicle control at 5hpi. Neutrophils at the wound site were photoconverted at 5hpi from green to red fluorescence. Time lapse imaging was performed from 7-12hpi. **B** The number of neutrophils which moved away from the wound site into a defined region of interest was quantified from 7-12 hours post injury in larvae treated with a vehicle control (black) or AMD3100 (red). Error bars shown are SEM, line of best fit shown is calculated by linear regression. P value shown is for the difference between the two slopes p<0.0001, n=35 larvae from 6 independent experiments. **C** Number of neutrophils photoconverted between 5-6 hours post injury in larvae treated with vehicle control or AMD3100. Data shown are mean ± SEM, groups were analysed using an unpaired t-test.

## Discussion

A large body of evidence now exists to suggest a role for the CXCL12/CXCR4 signalling axis in modulating neutrophil behaviour in chronic inflammatory disease. Aside from generation of neutrophil retention signals in multiple physiological settings [32], [33], neutrophils taken from patients with chronic inflammatory disease have increased CXCR4 expression, and CXCL12 is produced at sites of injury, including the lung [21], [22]. A specific role for the CXCL12/CXCR4 signalling axis in retaining neutrophils in the CHT has recently been suggested following the study of neutrophil behaviour in zebrafish Cxcr4b and Cxcl12a mutant larvae [30]. Our study provides evidence that the CXCL12/CXCR4 signalling axis is important in modulating neutrophil migration away from sites of inflammation, identifying a potential new therapeutic target for chronic inflammatory disease.

Computational modelling of reverse migration previously performed by our group demonstrated that neutrophil reverse migration is best described as a process of stochastic redistribution of neutrophils back into the tissue rather than their active migration away from the wound site [19]. These data further support our suggestion that neutrophil reverse migration is initiated following desensitisation to chemokine gradients at the wound site rather their active migration away from chemorepulsive gradients (fugetaxis). Cellular desensitisation to external gradients is a characteristic feature of signalling through G protein coupled receptors, many of which are expressed on the surface of neutrophils [34]. A retention signal generated through chemokine receptor signalling would require expression of the chemokine within the inflamed tissue and the receptor on the neutrophil surface. Our analysis of RNA sequencing from FACS sorted zebrafish larval neutrophils and adult single-cell RNA sequencing shows that at both larval and adult stages of development, the predominantly expressed isoform of CXCR4 in zebrafish neutrophils is *cxcr4b*, whilst *cxcr4a* was undetectable. This is in keeping with RT-PCR performed on FACS sorted larval zebrafish neutrophils [35]. Interestingly, RT-PCR performed on adult zebrafish whole kidney marrow suggests that both *cxcr4b* and *cxcr4a* are expressed by neutrophils in the adult stage [35]. Our analysis of single-cell RNA sequencing data provides a more sensitive assay to look at individual neutrophil RNA expression, therefore it is likely that zebrafish neutrophils do not express *cxcr4a* in adulthood. Furthermore, we demonstrate that mRNA for the major ligand for this receptor, Cxcl12a, is expressed at the wound site during inflammation. The *cxcl12a* expression pattern we observed in uninjured larvae was comparable to that observed by other groups earlier in zebrafish development at 2dpf [35]. Expression of *cxcl12a* mRNA appeared to increase at the wound site throughout the time course of inflammation, in keeping with a significant body of evidence that illustrates a role for CXCL12 in tissue repair [36]–[38]. It has been proposed that Cxcl12a is important in providing directional guidance cues to regulate endothelial cell migration during arterial morphogenesis in the regenerating fin [39]. Expression of *cxcl12a* is detected by WISH in injured adult tail fins from 1 day post amputation and persists during fin regeneration until 5 days post amputation [38].

The role for the CXCL12/CXCR4 signalling axis in zebrafish developmental processes has been elucidated largely using genetic studies to knock down the genes encoding the CXCR4 and CXCL12 proteins [27], [40], [41]. The high efficiency of somatic mutation by CRISPR/Cas9 in injected F0 animals yields up to 99% somatic mutagenesis and biallelic gene disruption, enabling direct phenotypic analysis without the requirement for raising stable F2 adults [29], [42], [43]. When using CRISPR/Cas9 to disrupt *cxcr4b* and *cxcl12a*, we achieved genomic disruption by introducing INDELs in >90% injected F0 larvae (Supplemental Figure 2). In our studies, knockdown of *cxcr4b* increased neutrophil recruitment to the wound site in crispant larvae. C-terminal truncations of Cxcr4b specifically in neutrophils (such as those found in WHIM syndrome patients) prevents receptor internalisation and increases sensitivity to Cxcl12a gradients, thus retaining them in the caudal hematopoietic tissue (CHT) inappropriately [35]. Neutrophils in WHIM zebrafish larvae are unable to respond to wound-generated gradients effectively, hence neutrophil recruitment to inflammatory sites is reduced in these larvae [35]. Conversely, in the Cxcr4b *odysseus* mutant where Cxcr4b signalling is impaired, the number of neutrophils available to be recruited to tissue damage is increased [30], thus our findings are in keeping with the F2 mutant phenotype [30]. Neutrophil recruitment towards Cxcl12a was not increased in our experiments, although this could be attributed to Cxcl12a larvae displaying significantly reduced whole body neutrophil counts. Inflammation resolution was significantly increased in both Cxcr4b and Cxcl12a crispant larvae, suggesting that genetic manipulation of both genes results in the same effect in terms of inflammation resolution.

One of the advantages of using the zebrafish as a model to study inflammation is that chemical compounds can be used to manipulate signalling pathways, where several compounds which target neutrophils have been identified using this approach [9], [11], [12]. AMD3100 is a non-peptide bicyclam which is able to specifically antagonize the CXCR4 receptor at three main interaction residues located around the main ligand binding pocket of CXCR4 in transmembrane domains IV, VI and VII. Binding of AMD3100 competitively inhibits binding of CXCL12 and prevents subsequent downstream signalling [44]. AMD3100 has been used to inhibit the CXCL12/CXCR4 signalling axis in zebrafish larvae, where concentrations ranging from 10-30µM have been administered to larvae through incubation in fish water for up to 24 hours [45], a concentration range which we remained within for our own experiments. Our results from both genetic and pharmacological manipulation of Cxcr4b and Cxcl12a demonstrate that inhibition of CXCL12/CXCR4 signalling accelerates inflammation resolution. We propose that AMD3100 is able to accelerate inflammation and reverse migration by competitively binding the CXCR4 receptor and preventing signalling downstream, thus recapitulating what would happen at a higher concentration of Cxcl12a later in the inflammatory response. AMD3100 can also act as an allosteric agonist of CXCR7 [46], which functions as a decoy receptor for CXCL12, with a role in cell generation of self-gradients which is crucial for proper migration of primordial germ cells toward their targets in zebrafish [47]. Activation of CXCR7 fails to couple to G-proteins and to induce chemokine receptor mediated cellular responses, so AMD3100 is unlikely to activate downstream signalling pathways [48]. Cxcr7 may modulate neutrophil sensitivity to Cxcl12, through its scavenging of the chemokine which reduces the level of Cxcl12 in the local tissue environment [49]. However, as zebrafish larval neutrophils do not express this receptor [25] (data not shown), it is unlikely that scavenging through Cxcr7 is involved.

Reverse migration is impaired in Cxcr2 deficient zebrafish larvae where neutrophils are inappropriately retained at the wound site [18]. It has been proposed that altered susceptibility of neutrophils to gradients at the wound site in Cxcr2 deficient larvae drives their passive migration away from the wound site. Our data are compatible with these findings, as the CXCR4 and CXCR2 signalling axis is known to antagonistically regulate neutrophil retention in other models [32]. It would be interesting to speculate that the combined outcome of signalling through both CXCR4 and CXCR2 could modulate the reverse migration of neutrophils during inflammation resolution.

Taken together our data demonstrate that inhibition of the CXCL12/CXCR4 signalling axis drives the resolution of inflammation by increasing neutrophil reverse migration, and supports the hypothesis that neutrophil desensitisation to gradients at the wound site results in their reverse migration away from the wound site [18], [19]. These data add to the existing evidence that neutrophil reverse migration can be targeted pharmacologically to drive the resolution of inflammation.

## Methods

### Zebrafish husbandry and ethics

To study neutrophils during inflammation *TgBAC(mpx:EGFP)i114* (known as mpx:GFP)[28] zebrafish larvae were in-crossed. To study gene expression by whole mount in situ hybridisation, wildtype pigment-less *nacre*[50] larvae were in-crossed. For reverse migration assays, *Tg(mpx:GAL4.vp16)sh267;Tg(UAS:Kaede)i222* (known as mpx:kaede) were in-crossed. All zebrafish were raised in the Bateson Centre at the University of Sheffield in UK Home Office approved aquaria and maintained following standard protocols[51]. Tanks were maintained at 28°C with a continuous re-circulating water supply and a daily light/dark cycle of 14/10 hours. All procedures were performed on embryos less than 5.2 dpf which were therefore outside of the Animals (Scientific Procedures) Act, to standards set by the UK Home Office.

### Neutrophil specific expression of zebrafish genes

Gene expression was assessed using an RNA sequencing database from FACS sorted GFP positive cells from 5dpf zebrafish and FPKM values for genes of interest were extracted [25] (data deposited on GEO under accession number GSE78954). For single cell analysis, gene expression values were extracted from the BASiCz (Blood atlas of single cells in zebrafish) cloud repository [26]. Cells of the neutrophil lineage were analysed for expression of *cxcr4a, cxcr4b, cxcl12a* and *cxcl12b.*

### WISH probe synthesis

The WISH antisense RNA probe for *cxcl12a* was synthesised from linearised plasmid DNA obtained from a plasmid vector containing the zebrafish *cxcl12a* coding sequence. Following transformation and DNA purification, the plasmid was linearised by restriction digest using EcoR1 (New England Biolabs (NEB), Herts, UK). The RNA probe was transcribed from linearised DNA using an SP6 RNA digoxigen labelling kit (Roche). 1μg of linearised DNA was incubated in a final volume of 20μl containing transcription reagents and transcription reaction was performed according to standard protocols (Roche).

### Whole mount in situ hybridisation

Nacre larvae were anaesthetised in tricaine following tail fin transection at time points indicated in the figure legends alongside uninjured, age-matched controls. No more than 20 larvae were transferred to 1ml Eppendorf tubes and excess liquid was removed without damaging larvae. 1ml of paraformaldehyde (PFA) at 4°c was added to Eppendorf tubes for the fixation step, and left overnight at 4°c. Larvae were washed and transferred into 100% methanol and stored at −20°c for at least 24 hours prior to use. WISH was performed using standard protocols [52] using an antisense DIG labelled probe for zebrafish *cxcl12a*.

### CRISPR/Cas9 reagents

Synthetic SygRNA® consisting of crRNA and tracrRNA (Merck) in combination with cas9 nuclease protein (Merck) was used for gene editing. Transactivating RNAs (tracrRNA) and gene specific CRISPR RNAs (crRNA) were resuspended to a concentration of 20µM in nuclease free water containing 10mM Tris-hcl ph8. SygRNA® complexes were assembled on ice immediately before injection using a 1:1:1 ratio of crRNA:tracrRNA:Cas9 protein. Gene-specific crRNAs to target *cxcr4b* and *cxcl12a* were designed using the online tool CHOPCHOP (http://chopchop.cbu.uib.no/). We used the following crRNA sequences, where the PAM site is indicated in brackets: ***cxcr4b:*** CAGCTCTGACTCCGGTTCTG(GGG) ***cxcl12a:*** CTCTACCAGGCTGATGGGCT(TGG).

### Microinjection of SygRNA® into embryos

A 1nl drop of SygRNA®:Cas9 protein complex was injected into mpx:GFP embryos at the one-cell stage. Embryos were collected at the one cell stage and injected using non-filament glass capillary needles (Kwik-Fil^TM^ Borosilicate Glass Capillaries, World Precision Instruments (WPI), Herts, UK). RNA was prepared in sterile Eppendorf tubes. A graticule was used to measure 0.5nl droplet sizes to allow for consistency of injections. Injections were performed under a dissecting microscope attached to a microinjection rig (WPI) and a final volume of 1nl was injected.

### Genotyping of crispant larvae

To determine the efficiency of CRISPR/Cas9 to induce site-specific mutations in injected larvae, we used restriction digest assays (Supplemental figure 2). CRISPR guides were designed to target sequences containing restriction digest sites, such that when indels were introduced by DNA repair, the restriction site is disrupted. Genomic DNA was extracted from individual larvae at 2dpf. Larvae were placed individually in 0.2ml PCR tubes in 90µl 50mM NaOH and boiled at 95° for 20 minutes. 10µl Tris-HCL ph8 was added as a reaction buffer and mixed thoroughly. RT-PCR using Firepol® (Solis BioDyne) was used to amplify a 235bp region (for *cxcr4b*) and a 259bp region (for *cxcl12a*) around the PAM site. Gene specific primers were designed using the Primer 3 web tool (http://primer3.ut.ee/). Primer sequences were as follows: ***cxcrb4_fw** TCCCGTATACTGTAGGGAGGA **cxcr4b_rev** TTTTTGCATTTTGTTTTCTTG **cxcl12a_fw** TTCTCTGTGGGACTGTGTTGAC **cxcl12a_rev**TTCGAAAATTTGACCCAAAAGT.* Restriction enzyme digests were then performed using bsII at 55° for 2 hours (for *cxcr4b*) and bstXi (New England Biolabs) at 37° for 2 hours (for *cxcl12a*). Products were run using gel electrophoresis on a 2% gel.

### Inflammation assays in crispant larvae

To induce an inflammatory response, chorions of zebrafish larvae at 2dpf were removed using sterile laboratory tweezers and larvae were anaesthetised in Tricaine (0.168 mg/ml; Sigma-Aldrich) in E3 media and visualised under a dissecting microscope. Tail-fins were transected consistently using a scalpel blade (5mm depth, WPI) by slicing immediately posterior to the circulatory loop, ensuring the circulatory loop remained intact as previously described[28]. Larvae were maintained at 28°c in fresh E3 media in a 24 well plate. Neutrophils at the wound site were counted at timepoints indicated in figure legends using a fluorescence stereo microscope.

### Compound treatment of larvae for inflammation resolution assays

To study the resolution of inflammation, neutrophils were counted at the wound site at intervals during the resolution phase from 8-24 hours post injury in 2dpf mpx:GFP larvae, as indicated in figure legends. Larvae were dechorionated and anaesthetised prior to injury by tail-fin transection and left to recover at 28°c in fresh E3 media in petri dishes (60 larvae per plate). Larvae were screened for good neutrophil recruitment (around 20 neutrophils at the wound site) at 3.5hpi. AMD3100 (Sigma-aldrich) was administered to larvae at 4hpi through injection into the duct of Cuvier at a final concentration of 20µM. AMD3100 was always tested alongside the appropriate vehicle control. Neutrophils at the wound site were counted at 6hpi at the peak of recruitment, and at 8hpi for inflammation resolution using a fluorescence stereo microscope (Leica).

### Percentage resolution calculations

To determine percentage resolution, experiments were performed with larvae maintained separately in a 96 well plate to follow individual larvae over time. Percentage resolution was calculated as ((Neutrophil counts at peak recruitment – neutrophil counts at 8hpi)/neutrophil counts at peak recruitment)*100.

### Whole body neutrophil counts

Whole body neutrophil counts were measured in mpx:GFP larvae at time points indicated in figure legends. Larvae were mounted in 1% agarose with tricaine and a single slice image was taken using a 4x NA objective lens on an Eclipse TE2000 U inverted compound fluorescence microscope (Nikon UK Ltd., Kingston upon Thames, UK). A GFP-filter was used at excitation of 488nm. Two images were taken per larvae, one of the head region and one of the tail region. Neutrophils were counted manually from both images and combined to give a whole body neutrophil count.

### Reverse migration assay

Reverse migration assays were performed using larvae expressing the photoconvertible protein kaede under the neutrophil specific mpx promoter: *TgBAC(mpx:GAL4-VP16); Tg(UAS:Kaede)i222*. At 3dpf larvae were anaesthetised and injured by tail-fin transection and left to recover at 28°c. Larvae were screened for good neutrophil recruitment at 4hpi. AMD3100 was administered by incubation in low melting point agarose containing tricaine at 5hpi in 3dpf larvae. Photoconverstion of kaede labelled neutrophils at the wound site was ^TM^ performed using an UltraVIEWPhotoKinesis device (Perkin Elmer and Analytical Sciences) on an UltraVIEWVoX spinning disc confocal laser imaging system (Perkin Elmer). The photokinesis device was calibrated using a coverslip covered in photobleachable substrate (Stabilo Boss^TM^, Berks UK). Photoconverstion was perfomed using a 405nm laser at 40% using 120 cycles, 250 pk cyles and 100ms as previously published [13]. Following calibration, a region of interest was drawn at the wound site between the edge of the circulatory loop and encapsulating the entirety of the wound edge. Successful photoconversion was detected through loss of emission detected following excitation at 488nm, and gain of emission following 561nm excitation. Larvae were then transferred to an Eclipse TE2000-U inverted compound fluorescence microscope with 10x NA objective lense to acquire images using an andor zyla 5 camera (Nikon). Time lapse imaging of neutrophil reverse migration was performed for 5 hours using 2.5 minute intervals using GFP and mCherry filters with 488 and 561 nm excitation respectively. For quantification of reverse migration, NIS elements software was used to compress z-slices into maximum intensity projections. A region of interest was drawn around the region away from the wound site, as illustrated in Figure 2.6. For quantification of neutrophils moving away from the wound site, a binary threshold was applied to images to detect mCherry neutrophils from background noise and NIS elements software calculated the number of objects detected in the ROI at each time point, providing a read out of reverse migration.

## Author Contributions

H.M.I performed all experiments with assistance from C.A.L, A.L.R and P.M.E. H.M.I and K.D.H analysed data. S.A.R and P.M.E conceived the study and designed experiments. S.A.R, P.M.E and L.R.P provided scientific expert knowledge. H.M.I wrote the manuscript with significant input from all authors.

## Acknowledgements

The authors would like to thank The Bateson Centre Aquarium Team at the University of Sheffield for their assistance with zebrafish husbandry. Imaging work was performed at the Wolfson Light Microscopy Facility, microscopy studies were supported by an MRC grant (G0700091) and a Wellcome Trust grant (GR077544AIA). Thanks to Dr Felix Ellett for use of zebrafish drawings for experiment schematics.

The authors declare no conflict of interest.

---

**Supplemental Figure 1.**
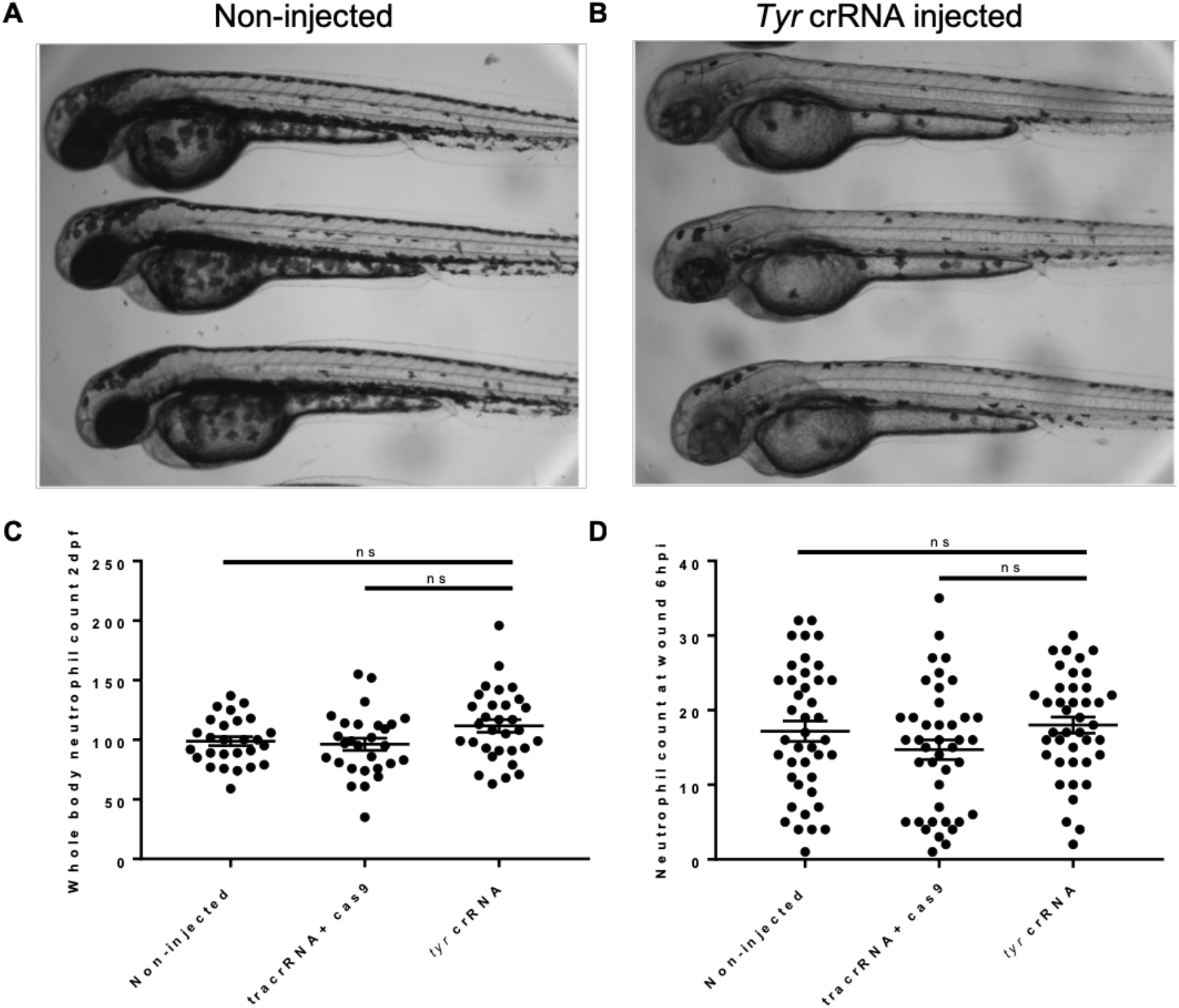
CRISPR/Cas9 knockdown of *tyrosinase* does not affect neutrophil function. **A-B.** Representative images of 2dpf mpx:GFP non-injected (**A**) and *tyrosinase* (**B**) mosaic pigment phenotypes. **C.** Whole body neutrophil counts in non-injected, vehicle control tracrRNA + cas9 protein injected and *tyrosinase* crRNA injected larvae. **D.** Neutrophils recruited to the injury site at 6hpi in 2dpf non-injected, vehicle control tracrRNA + cas9 protein injected and *tyrosinase* crRNA injected larvae. *(Error bars shown are mean ± SEM. Groups were analysed using an ordinary one-way ANOVA and adjusted using Tukeys multi comparison test, n=30 from 3 independent repeats*).

**Supplemental figure 2.**
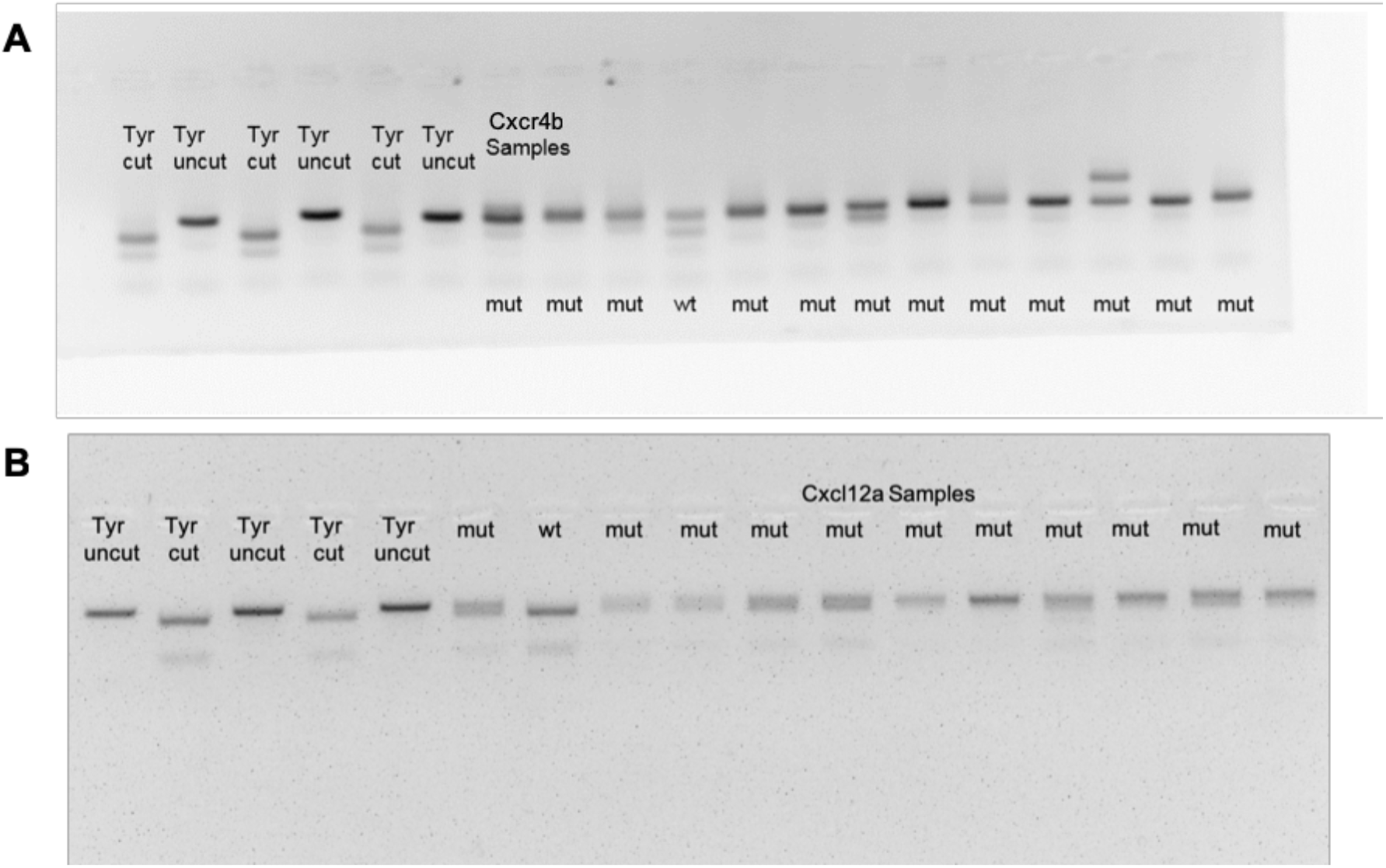
Genotyping of *cxcr4b* and *cxcl12a* CRISPR knockdown using restriction digest. **A** Electrophoresis gel for *cxcr4b* crispants at 2dpf. Lanes 1-6 Control *Tyr* injected larvae. Lanes 1,3,5 PCR produced incubated with bsII restriction enzyme, lanes 2,4,6 undigested PCR product. Lanes 7-19 *cxcr4b* crRNA injected larvae where PCR product has been digested using bsII. **B** Electrophoresis gel for *cxcl12a* crispants at 2dpf. Lanes 1-5 Control *Tyr* injected larvae. Lanes 1,3,5 Undigested PCR product, lanes 2,4 PCR produced incubated with bstXi restriction enzyme. Lanes 6-18 *cxcl12a* crRNA injected larvae where PCR product has been digested using bstXi.

## References

[1] C. Nathan and A. Ding, “Nonresolving Inflammation,” Cell, vol. 140, no. 6, pp. 871–882, Mar. 2010.

[2] T. A. Fuchs et al., “Novel cell death program leads to neutrophil extracellular traps,” J. Cell Biol., vol. 176, no. 2, pp. 231–241, Jan. 2007.

[3] C. Summers, S. M. Rankin, A. M. Condliffe, N. Singh, a. M. Peters, and E. R. Chilvers, “Neutrophil kinetics in health and disease,” Trends Immunol., vol. 31, no. 8, pp. 318–324, Aug. 2010.

[4] M. Mittal, M. R. Siddiqui, K. Tran, S. P. Reddy, and A. B. Malik, “Reactive Oxygen Species in Inflammation and Tissue Injury,” Antioxid. Redox Signal., vol. 20, no. 7, pp. 1126–1167, Mar. 2014.

[5] K. L. Rock, E. Latz, F. Ontiveros, and H. Kono, “The sterile inflammatory response.,” Annu. Rev. Immunol., vol. 28, pp. 321–42, 2010.

[6] Z. Bian, Y. Guo, B. Ha, K. Zen, and Y. Liu, “Regulation of the inflammatory response: enhancing neutrophil infiltration under chronic inflammatory conditions.,” J. Immunol., vol. 188, no. 2, pp. 844–53, Jan. 2012.

[7] J. M. Grigg, M. Silverman, J. S. Savill, C. Sarraf, and C. Haslett, “Neutrophil apoptosis and clearance from neonatal lungs,” Lancet, vol. 338, no. 8769, pp. 720–722, Sep. 1991.

[8] G. Cox, J. Crossley, and Z. Xing, “Macrophage engulfment of apoptotic neutrophils contributes to the resolution of acute pulmonary inflammation in vivo.,” Am. J. Respir. Cell Mol. Biol., vol. 12, no. 2, pp. 232–237, Feb. 1995.

[9] A. E. Leitch et al., “The cyclin-dependent kinase inhibitor R-roscovitine down-regulates Mcl-1 to override pro-inflammatory signalling and drive neutrophil apoptosis,” Eur. J. Immunol., vol. 40, no. 4, pp. 1127–1138, Apr. 2010.

[10] C. A. Loynes et al., “PGE _2_ production at sites of tissue injury promotes an anti-inflammatory neutrophil phenotype and determines the outcome of inflammation resolution in vivo,” Sci. Adv., vol. 4, no. 9, p. eaar8320, Sep. 2018.

[11] A. L. Robertson et al., “Identification of benzopyrone as a common structural feature in compounds with anti-inflammatory activity in a zebrafish phenotypic screen.,” Dis. Model. Mech., vol. 9, no. 6, pp. 621–32, 2016.

[12] A. L. Robertson et al., “A zebrafish compound screen reveals modulation of neutrophil reverse migration as an anti-inflammatory mechanism.,” Sci. Transl. Med., vol. 6, no. 225, p. 225ra29, 2014.

[13] P. M. Elks et al., “Activation of hypoxia-inducible factor-1?? (hif-1??) delays inflammation resolution by reducing neutrophil apoptosis and reverse migration in a zebrafish inflammation model,” Blood, vol. 118, no. 3, pp. 712–722, 2011.

[14] A. Woodfin et al., “The junctional adhesion molecule JAM-C regulates polarized transendothelial migration of neutrophils in vivo,” Nat. Immunol., vol. 12, no. 8, pp. 761–769, Aug. 2011.

[15] J. Wang, M. Hossain, A. Thanabalasuriar, M. Gunzer, C. Meininger, and P. Kubes, “Visualizing the function and fate of neutrophils in sterile injury and repair,” Science (80-.)., vol. 358, no. 6359, pp. 111–116, Oct. 2017.

[16] A. A. R. Thompson et al., “Hypoxia-inducible factor 2α regulates key neutrophil functions in humans, mice, and zebrafish.,” Blood, vol. 123, no. 3, pp. 366–76, Jan. 2014.

[17] K. Ley, C. Laudanna, M. I. Cybulsky, and S. Nourshargh, “Getting to the site of inflammation: the leukocyte adhesion cascade updated,” Nat. Rev. Immunol., vol. 7, no. 9, pp. 678–689, Sep. 2007.

[18] D. Powell, S. Tauzin, L. E. Hind, Q. Deng, D. J. Beebe, and A. Huttenlocher, “Chemokine Signaling and the Regulation of Bidirectional Leukocyte Migration in Interstitial Tissues.,” Cell Rep., vol. 19, no. 8, pp. 1572–1585, May 2017.

[19] G. R. Holmes et al., “Repelled from the wound, or randomly dispersed? Reverse migration behaviour of neutrophils characterized by dynamic modelling.,” J. R. Soc. Interface, vol. 9, no. 77, pp. 3229–39, Dec. 2012.

[20] T. Kawai and H. L. Malech, “WHIM syndrome: congenital immune deficiency disease.,” Curr. Opin. Hematol., vol. 16, no. 1, pp. 20–6, Jan. 2009.

[21] D. Hartl et al., “Infiltrated neutrophils acquire novel chemokine receptor expression and chemokine responsiveness in chronic inflammatory lung diseases.,” J. Immunol., vol. 181, no. 11, pp. 8053–67, Dec. 2008.

[22] M. Yamada et al., “The increase in surface CXCR4 expression on lung extravascular neutrophils and its effects on neutrophils during endotoxin-induced lung injury,” Cell. Mol. Immunol., vol. 8, no. 4, pp. 305–314, Jul. 2011.

[23] J. M. Petty et al., “Pulmonary stromal-derived factor-1 expression and effect on neutrophil recruitment during acute lung injury.,” J. Immunol., vol. 178, no. 12, pp. 8148–57, Jun. 2007.

[24] S. W. Chong, A. Emelyanov, Z. Gong, and V. Korzh, “Expression pattern of two zebrafish genes, cxcr4a and cxcr4b,” Mech. Dev., vol. 109, no. 2, pp. 347–354, Dec. 2001.

[25] J. Rougeot et al., “RNAseq profiling of leukocyte populations in zebrafish larvae reveals a cxcl11 chemokine gene as a marker of macrophage polarization during mycobacterial infection,” Front. Immunol., vol. 10, p. 832, 2019.

[26] E. I. Athanasiadis, J. G. Botthof, H. Andres, L. Ferreira, P. Lio, and A. Cvejic, “Single-cell RNA-sequencing uncovers transcriptional states and fate decisions in haematopoiesis,” Nat. Commun., vol. 8, no. 1, p. 2045, Dec. 2017.

[27] E. Donà et al., “Directional tissue migration through a self-generated chemokine gradient.,” Nature, vol. 503, no. 7475, pp. 285–9, 2013.

[28] S. A. S. Renshaw, C. A. Loynes, D. M. I. Trushell, S. Elworthy, P. W. Ingham, and M. K. B. Whyte, “A transgenic zebrafish model of neutrophilic inflammation,” Blood…, vol. 108, no. 13, pp. 3976–3978, Dec. 2006.

[29] L.-E. Jao, S. R. Wente, and W. Chen, “Efficient multiplex biallelic zebrafish genome editing using a CRISPR nuclease system,” Proc. Natl. Acad. Sci., vol. 110, no. 34, pp. 13904–13909, Aug. 2013.

[30] S. Paredes-Zúñiga et al., “CXCL12a/CXCR4b acts to retain neutrophils in caudal hematopoietic tissue and to antagonize recruitment to an injury site in the zebrafish larva,” Immunogenetics, vol. 69, no. 5, pp. 341–349, 2017.

[31] F. Ellett, P. M. Elks, A. L. Robertson, N. V. Ogryzko, and S. a. Renshaw, “Defining the phenotype of neutrophils following reverse migration in zebrafish,” J. Leukoc. Biol., vol. 98, no. December, pp. 1–7, Dec. 2015.

[32] K. J. Eash, A. M. Greenbaum, P. K. Gopalan, and D. C. Link, “CXCR2 and CXCR4 antagonistically regulate neutrophil trafficking from murine bone marrow,” J. Clin. Invest., vol. 120, no. 7, pp. 2423–2431, Jul. 2010.

[33] B. T. Suratt et al., “Role of the CXCR4/SDF-1 chemokine axis in circulating neutrophil homeostasis,” Blood, vol. 104, no. 2, pp. 565–571, Jul. 2004.

[34] A. C. Magalhaes, H. Dunn, and S. S. G. Ferguson, “Regulation of GPCR activity, trafficking and localization by GPCR-interacting proteins,” Br. J. Pharmacol., vol. 165, no. 6, pp. 1717–1736, 2012.

[35] K. B. Walters, J. M. Green, J. C. Surfus, S. K. Yoo, and A. Huttenlocher, “Live imaging of neutrophil motility in a zebrafish model of WHIM syndrome,” Blood, vol. 116, no. 15, pp. 2803–2811, Oct. 2010.

[36] J. Itou et al., “Migration of cardiomyocytes is essential for heart regeneration in zebrafish,” Development, vol. 139, no. 22, pp. 4133–4142, Nov. 2012.

[37] M. Bouzaffour, P. Dufourcq, V. Lecaudey, P. Haas, and S. Vriz, “Fgf and Sdf-1 pathways interact during zebrafish fin regeneration.,” PLoS One, vol. 4, no. 6, p. e5824, Jun. 2009.

[38] P. Dufourcq and S. Vriz, “The chemokine SDF-1 regulates blastema formation during zebrafish fin regeneration,” Dev. Genes Evol., vol. 216, no. 10, pp. 635–639, Oct. 2006.

[39] C. Xu et al., “Arteries are formed by vein-derived endothelial tip cells,” Nat. Commun., vol. 5, p. 5758, Dec. 2014.

[40] G. Valentin, P. Haas, and D. Gilmour, “The chemokine SDF1a coordinates tissue migration through the spatially restricted activation of Cxcr7 and Cxcr4b,” Curr. Biol., vol. 17, no. 12, pp. 1026–1031, 2007.

[41] P. Haas and D. Gilmour, “Chemokine Signaling Mediates Self-Organizing Tissue Migration in the Zebrafish Lateral Line,” Dev. Cell, vol. 10, no. 5, pp. 673–680, 2006.

[42] A. Burger et al., “Maximizing mutagenesis with solubilized CRISPR-Cas9 ribonucleoprotein complexes,” Development, vol. 143, no. 11, pp. 2025–2037, Jun. 2016.

[43] C. Cornet, V. Di Donato, and J. Terriente, “Combining Zebrafish and CRISPR/Cas9: Toward a More Efficient Drug Discovery Pipeline.,” Front. Pharmacol., vol. 9, p. 703, 2018.

[44] S. P. Fricker et al., “Characterization of the molecular pharmacology of AMD3100: A specific antagonist of the G-protein coupled chemokine receptor, CXCR4,” Biochem. Pharmacol., vol. 72, no. 5, pp. 588–596, 2006.

[45] O. J. Tamplin et al., “Hematopoietic stem cell arrival triggers dynamic remodeling of the perivascular niche,” Cell, vol. 160, no. 1–2, pp. 241–252, 2015.

[46] I. Kalatskaya, Y. A. Berchiche, S. Gravel, B. J. Limberg, J. S. Rosenbaum, and N. Heveker, “AMD3100 Is a CXCR7 Ligand with Allosteric Agonist Properties,” Mol. Pharmacol., vol. 75, no. 5, pp. 1240–1247, May 2009.

[47] B. Boldajipour et al., “Control of Chemokine-Guided Cell Migration by Ligand Sequestration,” Cell, vol. 132, no. 3, pp. 463–473, Feb. 2008.

[48] U. Naumann et al., “CXCR7 Functions as a Scavenger for CXCL12 and CXCL11,” PLoS One, vol. 5, no. 2, p. e9175, Feb. 2010.

[49] J. Bussmann and E. Raz, “Chemokine-guided cell migration and motility in zebrafish development,” EMBO J., vol. 34, no. 10, p. 1309, 2015.

[50] J. A. Lister, C. P. Robertson, T. Lepage, S. L. Johnson, and D. W. Raible, “nacre encodes a zebrafish microphthalmia-related protein that regulates neural-crest-derived pigment cell fate.,” Development, vol. 126, no. 17, pp. 3757–67, Sep. 1999.

[51] C. Nüsslein-Volhard and R. Dahm, Zebrafish : a practical approach. Oxford University Press, 2002.

[52] C. Thisse and B. Thisse, “High-resolution in situ hybridization to whole-mount zebrafish embryos.,” Nat. Protoc., vol. 3, no. 1, pp. 59–69, Jan. 2008.

